# Biocomputational Analysis Establishes Genetic Association of Rheumatoid Arthritis (RA) and Migraine

**DOI:** 10.1101/2020.02.05.936534

**Authors:** Nilanjan Roy, Arpita Mazumder

## Abstract

An autoimmune disease and a neurological disease do not tie up together but if statistics say that 61% of people are affected by Rheumatoid Arthritis (an autoimmune disease) and Migraine (a neurological disease) at the same time it is really a point to be noted. The previous studies show that Rheumatoid Arthritis (RA) and Migraine are related anatomically but our goal was to dig out any other similarities except it. Therefore, we studied their genes and hoped to get something fascinating. We assessed the genes from the literature study and Genetic Home Reference (https://ghr.nlm.nih.gov/). After that, the genetic sequences of these genes were extracted from NCBI (National Center for Biotechnology Information, http://ncbi.nlm.nih.gov) and by applying multiple sequence alignment, we created a phylogenetic tree by Mega (Molecular Evolutionary Genetics Analysis). Surprisingly, we found 10 pairs of genes with similar genetic structures and common ancestry. Therefore, we can say these 10 pairs of genes are related to each other closely, which is why Migraine and Rheumatoid Arthritis (RA) are found together in a person. At the last part of this study, we used protein expression for the evaluation of the result. Here our bioinformatics approach may help to strengthen the connectedness of these two diverse diseases.

## INTRODUCTION

A population-based study previously shown that there is a temporal relationship between Rheumatoid Arthritis (RA) and Migraine. Rheumatoid Arthritis (RA) is caused by an abnormality in the immune system that primarily affects joints by reducing synovial fluids. Variety of body systems, including the skin, eyes, lungs, heart, and blood vessels are also affected by Rheumatoid Arthritis (RA). Pain, swelling, stiffness, and loss of function in joints are common symptoms for Rheumatoid Arthritis (RA). About 2.1 million people, or between 0.5% and 1% of the U.S. adult population, have Rheumatoid Arthritis (RA). It is present in all races and age groups and occurs most frequently in women, 75% to be exact. Genetical and environmental reasons are suspected to cause Rheumatoid Arthritis (RA). Scientists have proved a specific genetic marker called the HLA (Human leukocyte antigen) shared epitope have a fivefold greater chance of developing Rheumatoid Arthritis (RA). The HLA genetic site controls immune responses. The role of HLA DRB1 (Major histocompatibility complex, class II, DR beta 1) alleles as a risk factor of RA has been known for 25 years. Associations between different HLA DRB1 alleles have been demonstrated in several populations across the world. There do appear to be differences in the strength of association between different alleles. For example, HLA DRB1*0404 is more commanding factor than HLA DRB1*0101. Furthermore, the full HLA DRB1 genotype is responsible for the risk of this disease not just the presence of one single allele. Individuals who carry the so-called ‘compound heterozygote genotype HLA DRB*0401/*0404 thus have a substantially greater risk than, for example, individuals who carry single HLA DRB*0101 alleles.^1^ PTPN22 (Protein tyrosine phosphatase, non-receptor type 22) is the first locus outside the HLA to demonstrate association with RA. PTPN22 is recognized as the second most important risk loci for Rheumatoid Arthritis (RA).^2^ The PTPN22 gene is a compelling biological candidate being a key component in the regulation of T cell receptor signaling.^3^ The associated risk allele encodes an arginine to tryptophan substitution at residue 620 in the polypeptide chain disrupting the binding to Csk. There is evidence that this change confers increased function to the PTPN22 protein, with the 620W variant enhancing the inhibitory effect on T cell receptor signaling.^4^ Another important locus related to Rheumatoid Arthritis (RA) is STAT4 (Signal transducer and activator of transcription 4). A combination of linkage and candidate gene studies showed the association of STAT4 with RA. In contrast with HLA-DRB1 and PTPN22, the association of STAT4 with RA is more modest.^2^ STAT4 is part of a family of seven transcription factors involved in cytokine receptor signaling, initially activated by IL-12 signaling in T cells, and then phosphorylated by janus kinases. It is also seen that STAT4 is translocated to the cell nucleus where it initiates transcription of its target genes and results in the production of interferon-γ and Th1 responses.^5^ Other genes connected to RA include AFF3, ARID5B, BLK C5, CCL21, CCR6, CD2, CD5, CD28, CD40, CD58, CTLA4, FCGR2A, FCGR2B, GATA3, IKZF3, IL2, IL2RA, IL2RB, IL6R, IL6ST, IL21, IRAK1, IRF5, IRF8, KIF5A, NFKBIL1, PADI4, PIP4K2C, POU3F1, PRDM1, PRKCQ, PTPRC, PXK, RASGRP1, RBPJ, RCAN1, REL, RUNX1, SPRED2, TAGAP, TLE3, TNFAIP3, TNFRSF14, TRAF1, TRAF6, TYK2.

Whereas a Migraine headache is usually an intense, throbbing pain on one, or sometimes, both sides of the head. In the ranks of most common disorder of the world, Migraine is the third most common affecting about one in seven people whereas about one in fifty people in chronic Migraine worldwide. Women tend to be affected by Migraines more frequently than men. Genetic, environmental, and lifestyle factors are the main factors to develop Migraine. Familial hemiplegic Migraine and Sporadic hemiplegic Migraine are most common in types. Familial hemiplegic Migraine is a form of Migraine headache that runs in families and sporadic hemiplegic Migraine is a rare form of Migraine headache. Migraine is caused by mutations in the CACNA1A (Calcium voltage-gated channel subunit alpha1 A), ATP1A2 (ATPase Na+/K+ transporting subunit alpha 2) and SCN1A (Sodium voltage-gated channel alpha subunit 1) genes.^6^ The CACNA1A gene sends signals for making one part (the alpha-1 subunit) of a calcium channel called CaV2.1. This subunit forms the pore through which calcium ions can flow. That is why CaV2.1 channels play an essential role in communication between nerve cells called neurons in the brain. These channels also help in controlling the release of neurotransmitters. Neurotransmitters are the chemicals that are released from nerve cells ends and relay signals from one neuron to another. Researchers believe that CaV2.1 channels are also involved in the survival of neurons-the ability of these cells to change and adapt over time (plasticity). The ATP1A2 gene provides instructions for making one part (the alpha-2 subunit) of a protein which is known as Na+/K+ ATPase. This enzyme uses energy from a molecule called adenosine triphosphate (ATP) to transport charged atoms (ions) into and out of cells. Specifically, it pumps potassium ions (K+) into cells while pumping sodium ions (Na+) out of cells against the concentration gradient. The SCN1A gene transmits signals for making one part (the alpha subunit) of a sodium channel called NaV1.1. These sodium channels are mostly found in the brain, where they control the flow of sodium ions into cells. NaV1.1 channels are involved in transmitting signals from one neuron to another. Communication between neurons depends on chemicals called neurotransmitters, after releasing from one neuron that taken up by neighboring neurons. The flow of sodium ions through NaV1.1 channels help to determine when neurotransmitters will be released. Recent reports suggested that the PRRT2 (Proline rich transmembrane protein 2) gene might be the fourth most important Migraine gene. Other genes associated with Migraine include ASTN2, CARF, CFDP1, HPSE2, IGSF9B, KCNK5, MEF2D, MPPED2, MRVI1, NRP1, PHACTR1, PLCE1, PRDM16, RNF213, SLC24A3, SUGCT, YAP1. The initial theory of people suffering from Rheumatoid Arthritis (RA) and Migraine together is that RA can causes decrease range of movement in the neck or certain movements and pressure worsen the pain in the neck. Neck with the help of spinal cord directly connects with human brain to cause Migraine. Resolving of the headache is usually done by blocking a cervical structure or its nerve supply. Some cervical abnormalities can trigger Migraine by activating the spinal trigeminal nucleus. When the cervical nerves communicate with the trigeminal nerves leading to trigeminocervical complex, which may result in arthritic pain at the front of the head. After studying genes responsible for Rheumatoid Arthritis (RA) and Migraine, we evaluated gene sequence similarity, and genetic significance between RA and Migraine through a computational tree method called phylogeny. Our study will look for reasons why patients suffering from RA is diagnosed with Migraine frequently.

**TABLE 1:** List of genes involved in Rheumatoid Arthritis (RA)

| **Genes associated with Rheumatoid Arthritis (RA)** | **Accession ID of NCBI** | **Tissue types** |
| --- | --- | --- |
| AFF3 | NM_002285.3 | Appendix, tonsil |
| ARID5B | NM_001244638.2 | Expressed in all |
| BLK | NM_001715.3 | Appendix, lymph node, spleen, tonsil, small intestine |
| C5 | NM_001735.2 | Liver |
| CCL21 | EF064765.1 | Lymph node, tonsil, small intestine, thyroid gland |
| CCR6 | NM_004367.5 | Lymph node, appendix, spleen, tonsil, small intestine, testis |
| CD2 | NM_001767.5 | Lymph node, spleen, small intestine, thymus |
| CD5 | NM_014207.4 | Lymph node, appendix, spleen, tonsil, small intestine, thymus |
| CD28 | NM_001243078.1 | Lymph node, spleen, thymus, tonsil, small intestine, placenta |
| CD40 | NM_001322421.2 | Expressed in all |
| CD58 | NM_001779.3 | Expressed in all |
| CTLA4 | AF414120.1 | Appendix, lymph node, tonsil, Small intestine |
| FCGR2A | NM_021642.4 | Placenta, Lung, spleen |
| FCGR2B | AH005422.2 | Placenta, spleen |
| GATA3 | NM_002051.3 | Parathyroid gland, skin, breast, placenta, urinary bladder, seminal vesicles, thymus |
| HLA-B | D83043.1 | Expressed in all |
| HLA-DPB1 | NM_002121.6 | Expressed in all |
| HLA-DRB1 | AB698851.1 | Lung |
| IKZF3 | NM_001257411.2 | Lymph node, appendix, spleen, tonsil, thymus, small intestine, thymus |
| IL2 | AH002842.2 | Evidence at protein level |
| IL2RA | NM_000417.2 | Lymph node, appendix, spleen, small intestine, adipose tissue |
| IL2RB | CR456506.1 | Placenta, spleen, tonsil |
| IL6R | NM_181359.3 | Mixed, expressed in all |
| IL6ST | AB102802.1 | Expressed in all |
| IL21 | NM_021803.4 | Lymph node, tonsil, testis |
| IRAK1 | NM_001025243.2 | Expressed in all |
| IRF5 | KP843528.1 | Spleen, seminal vesicle |
| IRF8 | NM_002163.4 | Lymph node, spleen, tonsil, small intestine |
| KIF5A | NM_001354705.2 | Cerebral cortex, cerebellum, hippocampus, caudate, brain |
| NFKBIL1 | KY500486.2 | Mixed, expressed in all |
| PADI4 | NM_012387.3 | Bone marrow, Lung, spleen |
| PIP4K2C | XM_017019973.2 | Expressed in all |
| POU3F1 | NM_002699.4 | Skin, caudate |
| PRDM1 | NM_001198.4 | Esophagus, expressed in all |
| PRKCQ | NM_001242413.2 | Skeletal muscle, thymus |
| PTPN22 | NM_015967.7 | Bone marrow, spleen, small intestine, thymus |
| PTPRC | NM_080921.3 | Lung, spleen, small intestine, expressed in all |
| PXK | KF774204.1 | Expressed in all |
| RASGRP1 | NM_001306086.2 | Cerebral cortex, cerebellum, lymph node, thymus |
| RBPJ | NM_015874.4 | Expressed in all |
| RCAN1 | NM_203418.3 | Parathyroid gland, expressed in all |
| REL | NM_001291746.2 | Expressed in all |
| RUNX1 | XM_005261068.3 | Expressed in all |
| SPRED2 | AY299090.1 | Expressed in all |
| STAT4 | EU304793.1 | Testis, pituitary gland |
| TAGAP | NM_152133.2 | Bone marrow, endometrium, lung, spleen, thymus, tonsil, small intestine |
| TLE3 | NM_020908.3 | Expressed in all |
| TNFAIP3 | NM_001270507.2 | Expressed in all |
| TNFRSF14 | NM_003820.3 | Expressed in all |
| TRAF1 | NM_005658.5 | Lymph node |
| TRAF6 | NM_145803.3 | Expressed in all |
| TYK2 | NM_003331.5 | Expressed in all |

**TABLE 2:** List of genes involved in Migraine

| **Genes associated with Migraine** | **Accession ID of NCBI** | **Tissue types** |
| --- | --- | --- |
| CACNA1A | NM_023035.3 | Cerebellum, cerebral cortex |
| ATP1A2 | NM_000702.4 | Cerebral cortex, skeletal muscle, brain, caudate, cerebellum, heart muscle, hippocampus, retina |
| SCN1A | MF358480.1 | Caudate, cerebellum, cerebral cortex, hippocampus, hypothalamus, lung |
| ASTN2 | NM_014010.5 | Expressed in all |
| CARF | NM_001352677.2 | Expressed in all |
| CFDP1 | CR542111.1 | Expressed in all, Evidence at protein level |
| HPSE2 | XM_024448120.1 | Cervix, uterine, smooth muscle |
| IGSF9B | XM_024448402.1 | Cerebellum |
| KCNK5 | NM_003740.4 | Gallbladder, kidney, small intestine |
| MEF2D | NM_001271629.2 | Expressed in all |
| MPPED2 | NM_001145399.2 | Cervix, uterine, thyroid gland |
| MRVI1 | NM_001100163.2 | Expressed in all |
| NRP1 | NR_045259.1 | Expressed in all |
| PHACTR1 | NM_001322314.2 | Cerebral cortex |
| PLCE1 | NM_016341.4 | Retina |
| PRDM16 | NM_199454.3 | Thyroid gland, retina |
| RNF213 | NM_020954.4 | Expressed in all |
| SLC24A3 | NM_020689.4 | Mixed |
| SUGCT | XM_011515530.3 | Kidney, adrenal gland |
| YAP1 | NM_006106.5 | Mixed, Expressed in all |

## MATERIALS AND METHODS

Literature study and with the help of genetic home reference (https://ghr.nlm.nih.gov/) we identified 20 genes associated with Migraine and 52 other genes associated with Rheumatoid Arthritis (RA). Our objective was to find the relation between these 72 genes in total to check if there was any relation between Migraine and Rheumatoid Arthritis (RA) genetically. In order to achieve that the functional DNA sequences of these total 72 genes were collected in fasta format from NCBI (National Center for Biotechnology Information, http://ncbi.nlm.nih.gov). Then we utilized these 72 genes and their DNA sequences with the help of Mega (Molecular Evolutionary Genetics Analysis) software (https://www.megasoftware.net/).^7^ To analyze the DNA sequences of these genes first we conducted multiple sequence alignment by clustalw method. Multiple sequence alignment is a method to identify related genes or proteins to find the evolutionary relationships between genes. It also reveals shared patterns among functionally or structurally related genes. Whereas clustalw is a progressive method to perform multiple sequence alignment. Clustalw compares only a pair of sequences together at a time. Using the standard dynamic programming algorithm on each pair, it can calculate the (N*(N-1))/2 (N is a total number of sequences) distances between the sequence pairs. A distance matrix is obtained using the cluster algorithm to construct a guide tree. After obtaining the tree, it aligns the first node to the second node. After fixing the alignment, add another sequence or the third node. Then it iterates the step until all the sequences are perfectly aligned. The alignment is performed that has the highest alignment score if a sequence is aligned to a group or if there is alignment in between the two groups of sequences. If it helps to guide the alignment of sequence, a gap symbol in the alignment is replaced with a neutral character.^8^ Then based on the result of multiple sequence alignment we constructed the phylogenetic tree with the help of maximum likelihood method to visualize the relationships among these 72 genes as mentioned. A phylogenetic tree is a tree diagram used to graphically show evolutionary relationships among organisms, DNA and protein sequences. A strong method to construct phylogenetic tree is the maximum likelihood method. It is a simple hill-climbing algorithm which adjusts the tree topology and branch lengths simultaneously. The algorithm starts from an initial tree built which is conducted by a fast distance-based method to modify this tree to improve its likelihood at each iteration successfully. As this is a simultaneous adjustment method, topology and branch lengths require only a few iterations to sufficiently reach an optimum.^9^ In the last step, we evaluated our result by protein expression analysis using protein atlas (https://www.proteinatlas.org/).^10,11, 12^

## RESULT

Construction of phylogenetic tree showed a notable relationship between genes of Rheumatoid Arthritis (RA) and Migraine. Genes that have most similar structure and most common ancestry are CD28 (a) and CARF (m), CD58(a) and CFDP1(m), TAGAP (a) and PLCE1 (m), IL6R (a) and ATP1A2 (m), TNFAIP3 (a) and PHACTR1 (m), SPRED2 (a) and MRVI1 (m), IL2RB (a) and PRDM16(m), HLA-B (a) and SLC24A3(m), RUNX1 (a) and SUGCT (m), STAT4 (a) and MPPED2 (m) where (a) denotes for gene of Rheumatoid Arthritis (RA) and (m) denotes for gene of Migraine (figure: 1). This analysis of genes giving us result that genes responsible for Rheumatoid Arthritis (RA) and Migraine indeed have structural similarities and therefore related to each other in a manner that people having Rheumatoid Arthritis (RA) are susceptible to Migraine or people having Migraine are susceptible to Rheumatoid Arthritis (RA). Further protein expression analysis was done on top genes responsible for Rheumatoid Arthritis (RA) and Migraine. Top genes responsible for Rheumatoid Arthritis (RA) are HLA-B, PTPN22, and STAT4, for Migraine are CACNA1A, ATP1A2, and SCN1A. The phylogenetic tree showed us relationship between these top genes such as HLA-B (a) and SLC24A3 (m), STAT4 (a) and MPPED2 (m) and IL6R (a) and ATP1A2 (m). HLA-B, SLC24A3, STAT4, MPPED2, IL6R, ATP1A2 denotes for Major histocompatibility complex, class I, B, Solute carrier family 24 member 3, Signal transducer and activator of transcription 4, Metallophosphoesterase domain containing 2, Interleukin 6 receptor, ATPase Na+/K+ transporting subunit alpha 2 respectively.

**FIGURE 1:**
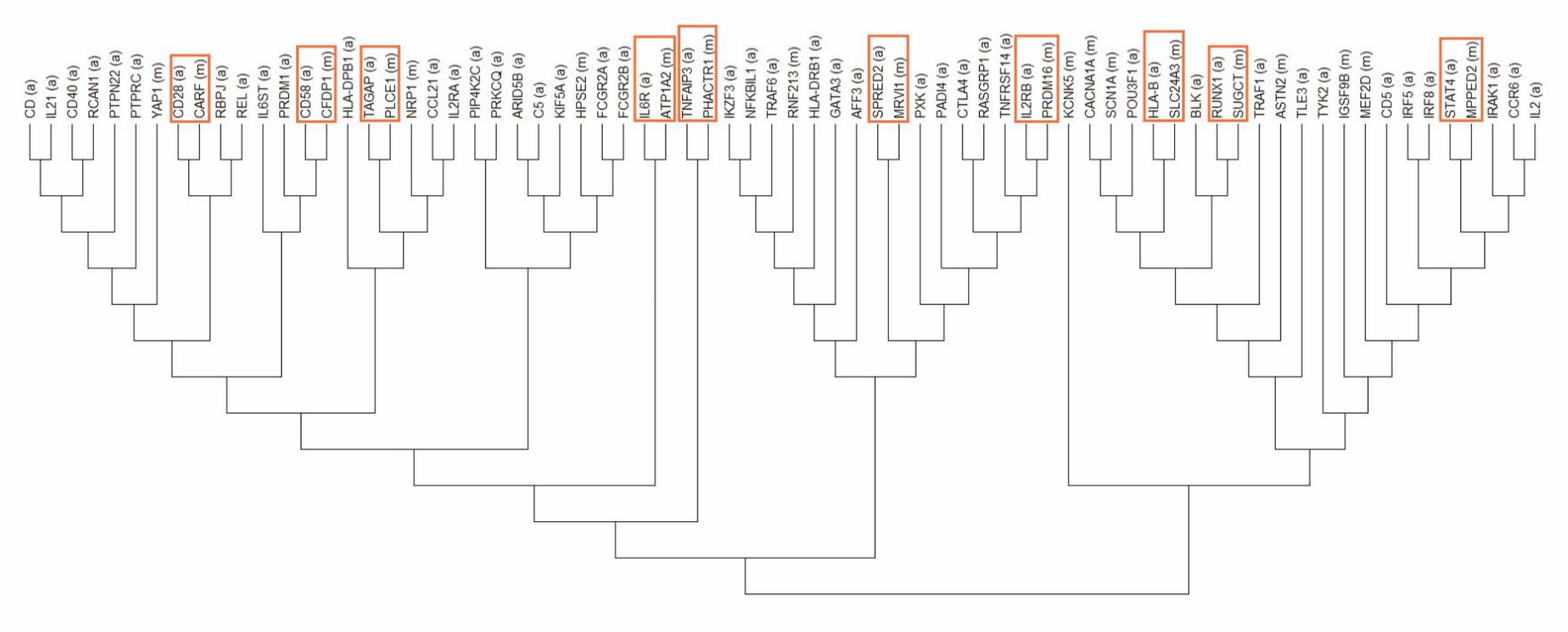
Shows the relationship between the genes by using the phylogenetic tree between Rheumatoid Arthritis (RA) and Migraine. (a) denotes as gene for Rheumatoid Arthritis (RA) and (m) denotes as gene for Migraine.

We know that Rheumatoid Arthritis (RA) is a disease of immune system disorder and in relation to that HLA-B gene which is the main cause of this disease shows high protein expression of bone marrow and spleen (figure: 2). Bone marrow and spleen produces T-cells and B-cells which makes up our body’s immune system. Whereas Migraine symptoms are situated in the nervous system or brain. Protein expression overview graph indicates that even HLA-B gene shows low protein expression in cerebral cortex. HLA-B gene is highly related with SLC24A3 gene of Migraine and this SLC24A3 gene shows medium level protein expression among cerebral cortex, cerebellum, and spleen (figure: 3). STAT4 is also an important gene to cause Rheumatoid Arthritis (RA). STAT4 shows low-level spleen, bone marrow, lymph node protein expression to cause Rheumatoid Arthritis (RA). On the other hand, it also shows medium level protein expression in the cerebral cortex and low-level protein expression in the cerebellum to cause Migraine (figure: 4). Another important gene to consider for Migraine is ATP1A2. It has high-level protein expression for cerebral cortex and medium level protein expression in cerebellum for developing Migraine (figure: 5). Besides that, ATP1A2 gene shows low-level skeletal muscle protein expression. Skeletal muscle is directly related to the disease Rheumatoid Arthritis (RA). Therefore, phylogenetic analysis for 72 genes and further evaluating protein expression of major genes responsible for Rheumatoid Arthritis (RA) and Migraine indicate strong associations of genes between these two diseases.

**FIGURE 2:**
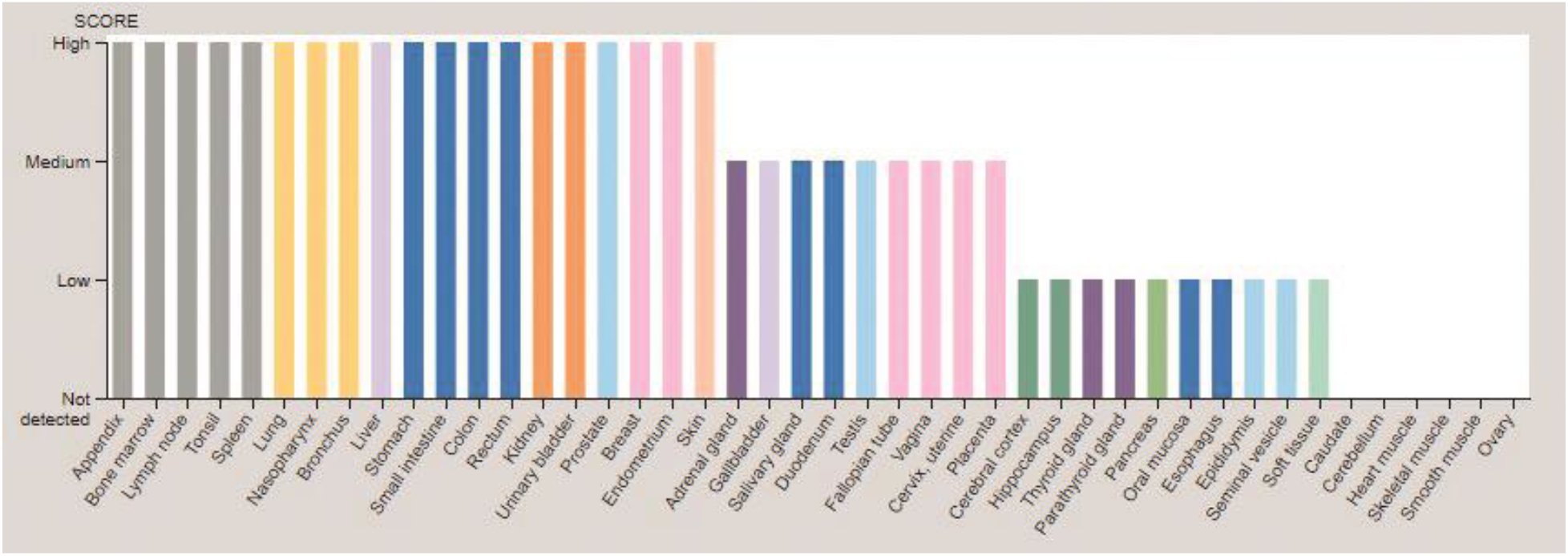
Protein expression overview of gene HLA-B that is responsible for Rheumatoid Arthritis (RA). Image credit: Human Protein Atlas (https://www.proteinatlas.org/).

**FIGURE 3:**
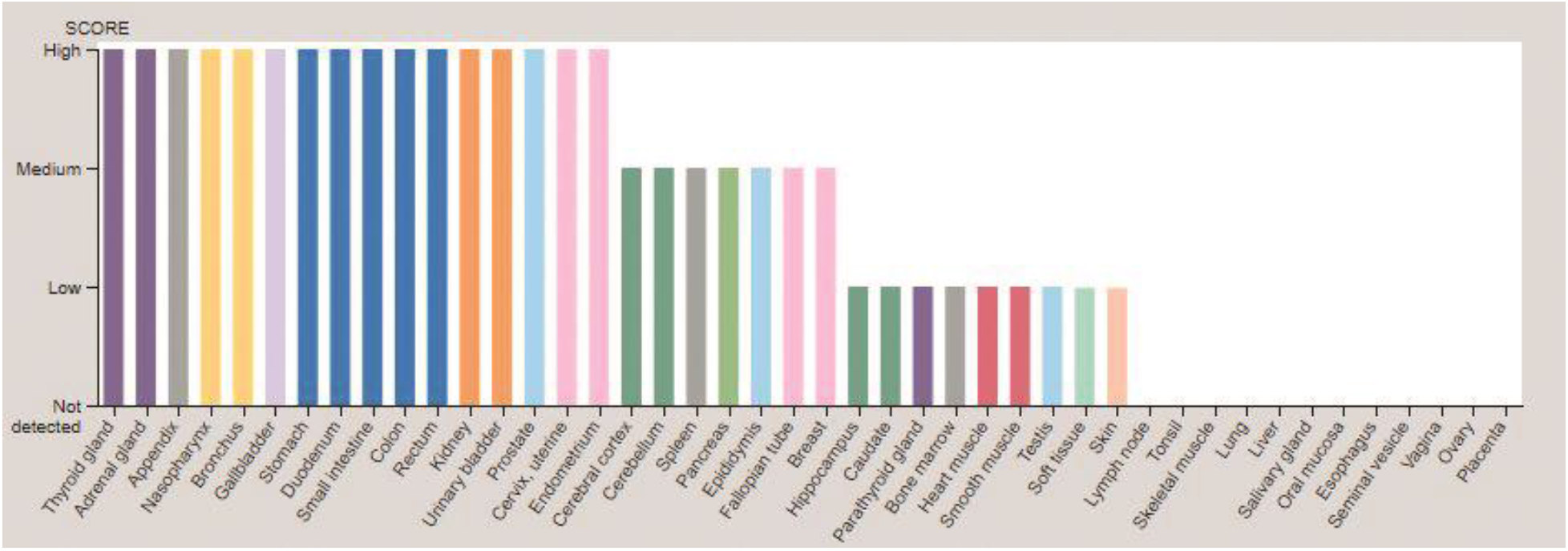
Protein expression overview of gene SLC24A3 that is responsible for Migraine. Image credit: Human Protein Atlas (https://www.proteinatlas.org/).

**FIGURE 4:**
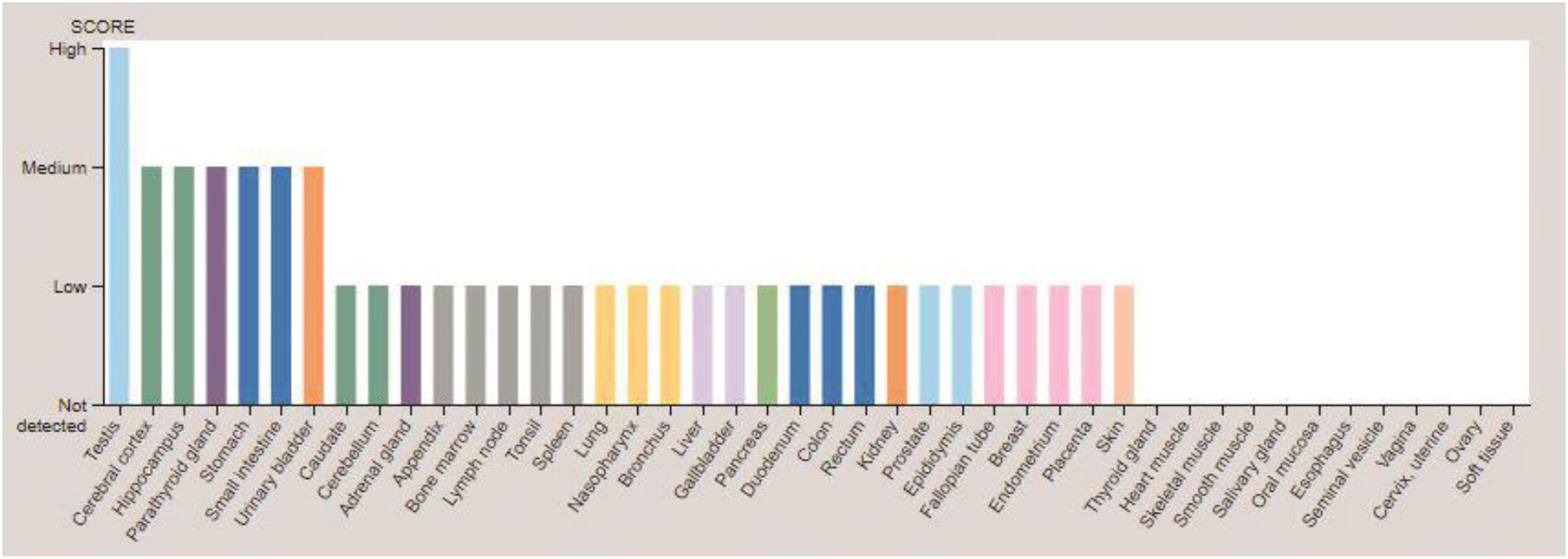
Protein expression overview of gene STAT4 that is responsible for Rheumatoid Arthritis (RA). Image credit: Human Protein Atlas (https://www.proteinatlas.org/).

**FIGURE 5:**
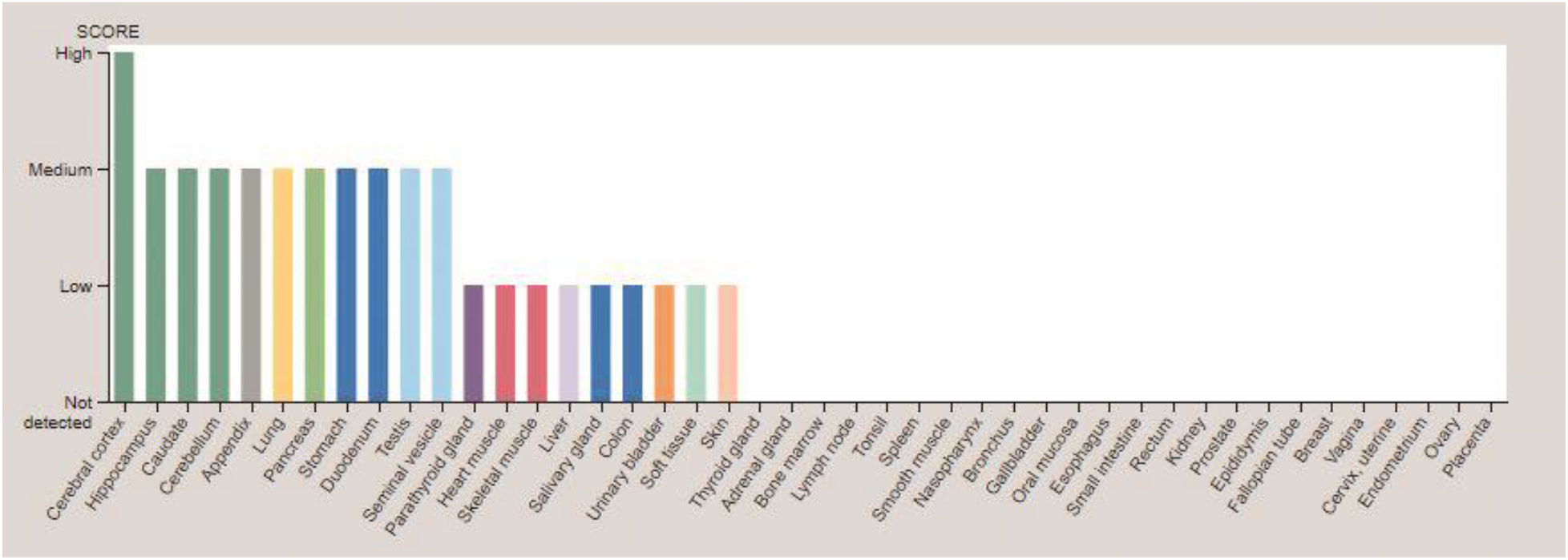
Protein expression overview of gene ATP1A2 that is responsible for Migraine. Image credit: Human Protein Atlas (https://www.proteinatlas.org/).

## DISCUSSION

Genetic similarity of Rheumatoid Arthritis (RA) and Migraine could be established which was the sole purpose of this study. The desired result is found as the 10 pairs of genes from the phylogenetic tree analysis have common ancestry and genetic similarity (figure: 1). Therefore, it can be said that these 10 pairs of genes are responsible to cause Rheumatoid Arthritis (RA) and Migraine at the same time in patients. However, all of these 20 genes are not the prime genes to cause the above diseases. So, in our results, we discussed only the chief three gene pairs and among them two are prime genes (HLA-B and STAT4) to cause Rheumatoid Arthritis (RA) and one (ATP1A2) is the leading gene that causes Migraine. The protein expression of these chief genes are analyzed from The Human Protein Atlas (https://www.proteinatlas.org/) to further strengthen our phylogenetic tree relation for evaluation of our result. Our study is based on the phylogenetic tree and we used maximum likelihood method. In this model, the assumptions are clear and statistically efficient to construct the perfect phylogenetic tree. A limitation of maximum likelihood is that it is computationally slower. Aside from that, the likelihood ratio test could not be used to test the tree topologies as it is not supported by the method, we used to build the phylogenetic tree. Therefore, it is another drawback of maximum likelihood method. On the other side, the protein expression of the top genes is drawn from The Human Protein Atlas (https://www.proteinatlas.org/) that is why the result’s validity is dependent on the efficacy of the protein atlas data undoubtedly.

## CONCLUSION

Rheumatoid Arthritis (RA) and Migraine both are chronic diseases so there is no cure but long-term sufferings. The curative and preventive procedures would be attainable if further investigation with more genes (both with the discovered genes and upcoming discoveries) and more methods is applied to analyze these genes. Our study was only the first step based on the theoretical framework and biocomputational power to find the association between Rheumatoid Arthritis (RA) and Migraine. Their parallelism in a person might be concluded one day if more detailed study is carried out to find more possible relativity.

## REFERENCES

1 Silman, A. and Pearson, J. (2002). Epidemiology and genetics of Rheumatoid Arthritis (RA). Arthritis Research, 4(Suppl 3), p.S265.

2 Orozco, G., McAllister, K. and Eyre, S. (2011). Genetics of Rheumatoid Arthritis (RA): GWAS and beyond. Open Access Rheumatology: Research and Reviews, p.31.

3 Bottini N, Musumeci L, Alonso A, et al. A functional variant of lymphoid tyrosine phosphatase is associated with type I diabetes. Nat Genet. 2004; 36(4):337–338.

4 Vang T, Congia M, Macis MD, et al. Autoimmune-associated lymphoid tyrosine phosphatase is a gain-of-function variant. Nat Genet. 2005; 37(12):1317–1319.

5 Wurster AL, Tanaka T, Grusby MJ. The biology of Stat4 and Stat6. Oncogene. 2000; 19(21):2577–2584.

6 De Boer, I., van den Maagdenberg, A. and Terwindt, G. (2019). Advance in genetics of Migraine. Current Opinion in Neurology, 32(3), pp.413–421.

7 Sudhir Kumar, Glen Stecher, Michael Li, Christina Knyaz, and Koichiro Tamura (2018) MEGA X: Molecular Evolutionary Genetics Analysis across computing platforms. Molecular Biology and Evolution 35:1547–1549.

8 Daugelaite, J., O’ Driscoll, A. and Sleator, R. (2013). An Overview of Multiple Sequence Alignments and Cloud Computing in Bioinformatics. ISRN Biomathematics, 2013, pp.1–14.

9 Guindon, S. and Gascuel, O. (2003). A Simple, Fast, and Accurate Algorithm to Estimate Large Phylogenies by Maximum Likelihood. Systematic Biology, 52(5), pp.696–704.

10 Uhlen, M., Fagerberg, L., Hallstrom, B., Lindskog, C., Oksvold, P., Mardinoglu, A., Sivertsson, A., Kampf, C., Sjostedt, E., Asplund, A., Olsson, I., Edlund, K., Lundberg, E., Navani, S., Szigyarto, C., Odeberg, J., Djureinovic, D., Takanen, J., Hober, S., Alm, T., Edqvist, P., Berling, H., Tegel, H., Mulder, J., Rockberg, J., Nilsson, P., Schwenk, J., Hamsten, M., von Feilitzen, K., Forsberg, M., Persson, L., Johansson, F., Zwahlen, M., von Heijne, G., Nielsen, J. and Ponten, F. (2015). Tissue-based map of the human proteome. Science, 347(6220), pp.1260419–1260419.

11 Thul, P., Åkesson, L., Wiking, M., Mahdessian, D., Geladaki, A., Ait Blal, H., Alm, T., Asplund, A., Björk, L., Breckels, L., Bäckström, A., Danielsson, F., Fagerberg, L., Fall, J., Gatto, L., Gnann, C., Hober, S., Hjelmare, M., Johansson, F., Lee, S., Lindskog, C., Mulder, J., Mulvey, C., Nilsson, P., Oksvold, P., Rockberg, J., Schutten, R., Schwenk, J., Sivertsson, Å., Sjöstedt, E., Skogs, M., Stadler, C., Sullivan, D., Tegel, H., Winsnes, C., Zhang, C., Zwahlen, M., Mardinoglu, A., Pontén, F., von Feilitzen, K., Lilley, K., Uhlén, M. and Lundberg, E. (2017). A subcellular map of the human proteome. Science, 356(6340), p.eaal3321.

12 Uhlen, M., Zhang, C., Lee, S., Sjöstedt, E., Fagerberg, L., Bidkhori, G., Benfeitas, R., Arif, M., Liu, Z., Edfors, F., Sanli, K., von Feilitzen, K., Oksvold, P., Lundberg, E., Hober, S., Nilsson, P., Mattsson, J., Schwenk, J., Brunnström, H., Glimelius, B., Sjöblom, T., Edqvist, P., Djureinovic, D., Micke, P., Lindskog, C., Mardinoglu, A. and Ponten, F. (2017). A pathology atlas of the human cancer transcriptome. Science, 357(6352), p.eaan2507.

